# Colonial Architecture Modulates the Speed and Efficiency of Multi-Jet Swimming in Salp Colonies

**DOI:** 10.1101/2024.04.18.590155

**Authors:** Alejandro Damian-Serrano, Kai A. Walton, Anneliese Bishop-Perdue, Sophie Bagoye, Kevin T. Du Clos, Bradford J. Gemmell, Sean P. Colin, John H. Costello, Kelly R. Sutherland

## Abstract

Salps are marine pelagic tunicates with a complex life cycle including a solitary and colonial stage. Salp colonies are composed of asexually budded individuals that coordinate their swimming by multi-jet propulsion. Colonies develop into species-specific architectures with distinct zooid orientations. We hypothesize that colonial architecture drives differences in swimming performance between salps due to differences in how frontal drag scales with the number of propeller zooids in the colony. Moreover, we hypothesize that faster-swimming taxa are more energetically efficient in their locomotion since less energy would be devoted to overcoming drag forces. We (1) compare swimming speed across salp species and architectures, (2) evaluate how swimming speed scales with the number of zooids in the colony in architectures with constant and scaling frontal cross-sectional area, and (3) compare the metabolic cost of transport across different species and how it scales with swimming speed. To measure their swimming speeds, we recorded swimming salp colonies using in situ videography while SCUBA diving in the open ocean. To estimate the cost of transport, we measured the respiration rates of swimming and anesthetized salps collected in situ using jars equipped with non-invasive oxygen sensors. We found that linear colonies generally swim faster and with a lower cost of transport due to their differential advantage in frontal drag scaling with an increasing number of zooids. These findings underscore the importance of considering propeller arrangement to optimize speed and energy efficiency in bioinspired underwater vehicle design, leveraging lessons learned from the diverse natural laboratory provided by salp diversity.

**Summary Statement:** Linear arrangements in multi-jet propelled marine colonial invertebrates are faster and more energetically efficient than less streamlined architectures, offering insights for bioinspired underwater vehicle design.

## Introduction

Salps (Tunicata: Thaliacea: Salpida) are planktonic invertebrates that have a two-phase life cycle comprised of a solitary oozooid that asexually buds colonies of sexually reproducing blastozooids. Salp colonies are composed of up to hundreds of genetically identical, physically and neurologically integrated pulsatile zooids (Bone et al. 1980, Mackie 1986). Zooids in the colony feed and propel themselves by inhaling water through the oral siphon, using muscle contraction to compress their pharyngeal chamber, and exhaling a jet of water from their atrial siphon (Bone & Trueman 1983). While solitary oozooids move using single-jet propulsion, salp blastozooid colonies integrate multiple propelling jets, which increases their thrust and reduces the drag that results from periodical acceleration and deceleration via asynchronous swimming (Sutherland & Weihs 2017).

Currently, there are 48 described species of salps (WoRMS, 2024) and differences between species have mainly been compared from a taxonomic lens, focused on zooid-level diagnostic morphological characters. While salps are widely distributed, most salp species are restricted to open ocean environments, far from the coast with extremely deep bottom depths, which poses unique challenges to accessing them for direct study in their environment (Hamner et al 1975, Haddock 2004). Moreover, salps cannot be maintained alive in containers beyond a few hours since they are extremely fragile and sensitive to the presence of solid walls. Therefore, many morphological, ecological, and functional aspects of salp diversity, such as swimming speeds and metabolic demands, have remained unexplored. One such aspect is colonial architecture or the way that the zooids are arranged relative to each other in the colony. Salp colonies develop into species-specific architectures with distinct zooid orientations, including transversal, oblique, linear, helical, and bipinnate chains; as well as whorls, and clusters (Damian-Serrano & Sutherland, 2023). These architectures present distinct orientations of the propeller zooids and their thrusting jets to the axes of colony elongation and locomotion hypothesized to have an impact on their swimming performance (Madin 1990, Damian-Serrano et al. 2023).

Linear salp chains have been hypothesized to be more efficient swimmers due to the reduction of drag associated with a more streamlined form (Bone & Trueman 1983). We expect frontal drag scaling to be relevant to the hydrodynamics of swimming salp colonies given that their intermediate Reynolds numbers are estimated to be between ∼100 (Sutherland & Madin 2010) for solitary salps and ∼5000 for linear chains (Sutherland & Weihs 2017). In animals swimming at high Reynolds numbers, such as colonial salps, the drag experienced during swimming depends largely on the frontal (motion-orthogonal) projected area (Alexander 1968, Vogel 1981). Having a larger number of propellers is expected to improve the hydrodynamic and inertial benefits granted by asynchronous multijet propulsion, in addition to providing additional thrust to the colony (Madin 1990, Sutherland & Weihs 2017). The effect of varying numbers of propeller zooids on swimming speed has never been investigated in salps, nor how this relationship may vary across their diverse colonial architectures. While relative frontal drag is greatly reduced in linear chains when compared to the sum of each separate blastozooid (Mackie 1986, Sutherland & Weihs 2017), we hypothesize that this advantage will be lower in species with non-linear colonial architectures, and thus we predict finding differences in swimming speed between colonial architectures. Salp colonial architectures differ in how the number of zooids in the colony scales with their frontal area relative to motion (Madin 1990). Some architectures (linear, bipinnate, and helical) have a constant frontal area relative to their motion, regardless of zooid number. We expect these architectures to benefit from increased thrust delivered by larger numbers of zooids while maintaining a constant frontal drag resistance. However, the rest of the architectures (oblique, transversal, whorl, and cluster) have an increasing frontal area as the number of zooids increases (Fig. 1). Therefore, we expect the latter architectures to not only obtain more thrust, but to also experience more frontal drag as a result of bearing a greater number of propeller zooids. As a result, we also predict that swimming speed will be greater in colonies that bear a larger number of zooids, but only for species with architectures that have a constant frontal area.

**Figure 1.**
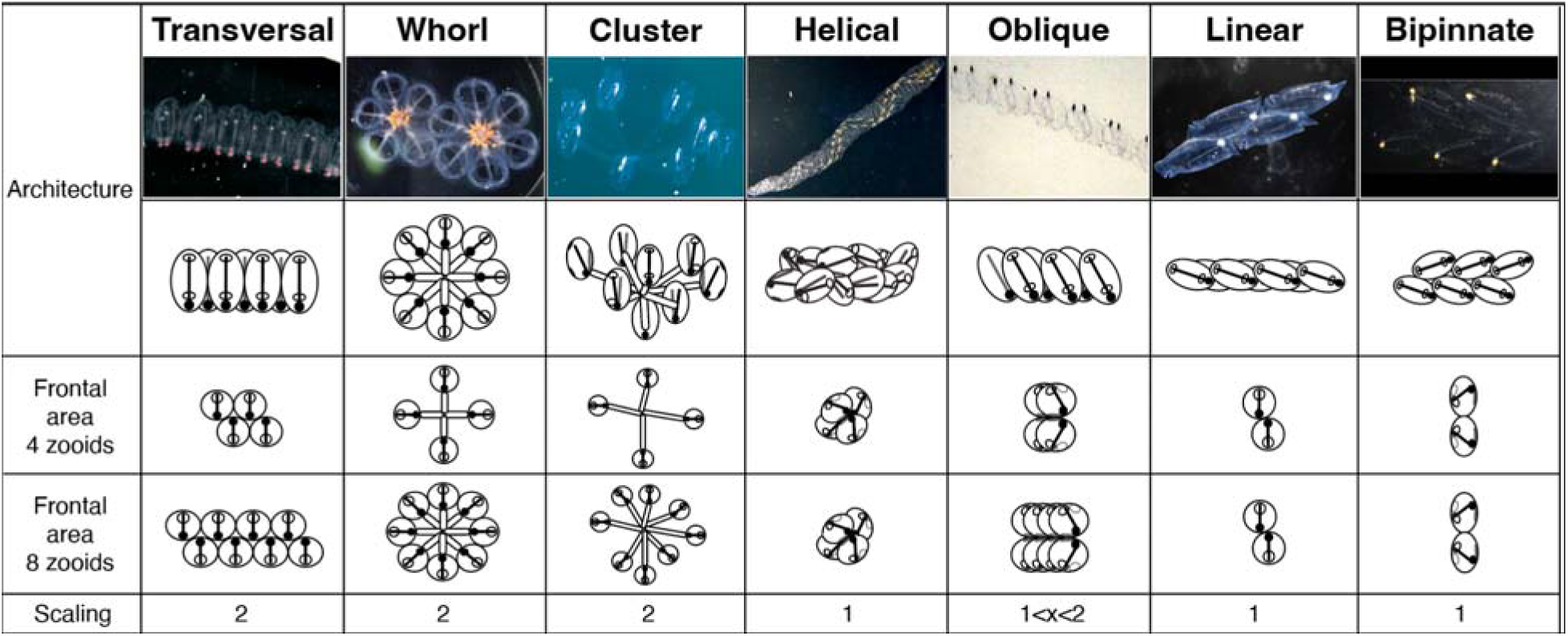
Salp colonial architectures with representative species photos (*Pegea* sp. For transversal, *Cyclosalpa affinis* for whorl, *Cyclosalpa sewelli* for cluster, *Helicosalpa virgula* for helical, *Thalia cicar* for oblique, *Soestia zonaria* for linear, and *Ritteriella retracta* for bipinnate) and diagrams showing the distinct zooid orientations. The subsequent rows show the frontal view of colonies with four and eight zooids, with the final row indicating the expected frontal area increase factor between the four and the eight zooid colonies.

Among the architectures with constant frontal area, we expect linear chains to be the fastest due to having the most streamlined arrangement of zooids parallel or near parallel to the axis of motion (Damian-Serrano & Sutherland, 2023), followed by the helical and bipinnate chains in which the zooids are angled relative to the axis of motion. Among the architectures with increasing frontal area with the number of zooids, we expect oblique chains to be the fastest, while still slower than those with constant frontal area, since their arrangement is partially aligned (angled dorsoventrally) with the axis of motion (Damian-Serrano & Sutherland 2023). Since both whorls and transversal chains have zooids rigidly attached at 90-degree angles to the axis of motion, we expect them both to have similar swimming speeds (slower than oblique) and scaling rates with the number of zooids in the colony. In cluster colonies the zooids are attached to a center point solely by their long flexible peduncle, which allows them to bend their orientation and pivot back and forth as a result of their jet propulsion. This may shunt thrust from propulsion into zooid-pivoting torque, thus we expect these colonies to be the slowest swimmers. Salp zooids pump water as a means of filter feeding as well as to move in the water column. The latter function is particularly relevant for species that undergo diel vertical migration, which not all species do (Madin et al. 1996). Therefore, the eco-evolutionary relevance of swimming speed and the hydrodynamic efficiency may vary between species (Damian-Serrano et al. 2023).

The degree of linearity in a colony can be expressed as the degree of parallelism between the zooids and the elongation axis of the chain. This angle is determined by the degree of developmental dorsoventral zooid rotation, which can span from 90°, in transversal chains with no rotation, to 0° (perfect linearity), in some linear chains such as those from the species *Soestia zonaria* (Damian-Serrano & Sutherland, 2023). Strong reductions in the dorsoventral zooid rotation angle toward linear forms have evolved multiple times independently (Damian-Serrano et al. 2023), possibly due to adaptive advantages related to their swimming efficiency. Therefore, based on the same rationale as for the abovementioned hypotheses, we further hypothesize that swimming speed is faster in species with lower dorsoventral zooid rotation angle.

Madin (1990) found a linear relationship between swimming effort (pulsation rate) and swimming velocity in solitary zooids. We hypothesize this relationship to also be present in colonial zooids. While body size predicts swimming velocity in many animals (Vogel 2008), Madin (1990) did not find such a relationship in salp blastozooids or oozooids. Since asynchronous-pulsating cruising salp colonies overcome many of the acceleration issues that limit single-jetters (Sutherland & Weihs 2017), we hypothesize that in salp colonies, zooid (propeller) size will be predictive of swimming speed across species.

The energetic costs of salp locomotion have been previously estimated using mechanically estimated propulsive efficiency as a proxy in three species (Sutherland & Madin 2010, Gemmell et al. 2021) and with a direct comparison between swimming and anesthetized respiration rates in *Salpa fusiformis* (Trueman et al. 1984). The metabolic demands of salp colonies have been estimated for a few species of salps in context with other gelatinous zooplankton (Biggs 1977, Schneider 1992, Mayzaud et al. 2005, Trueblood 2019), showing that salps have a relatively higher respiration rate than other gelatinous taxa. Cetta et al. (1986) compared the respiration rates across salp species to their pulsation rate and swimming speeds, revealing that more active species had higher respiration rates. However, the specific costs incurred by their swimming activity and their relationship to swimming speed have never been examined across the diversity of salp species. We hypothesize that species with a higher overall pulsation rate invest more of their metabolic demands in swimming. If faster swimming salp species are faster due to experiencing less frontal drag force as a result of their colonial architecture, we hypothesize that their swimming should also be less costly, since they would spend less energy in overcoming the forces opposing their forward motion. Under this hypothesis, we would predict that faster species will present lower costs of transport (energetic costs of displacement per unit of distance).

In this study, we compare the swimming speeds across 17 salp species and the energetic costs of swimming across 15 species of salps, encompassing all six different salp colony architectures. In addition, we investigate how swimming speed varies with the number of propeller zooids and evaluate whether differences in frontal area scaling drive disparities between colonial architectures. Finally, we assess how the cost of transport of salp colony swimming varies between species, as well as how their swimming efficiency scales with swimming speed and effort.

## Materials and Methods

### Fieldwork

We observed salps via bluewater SCUBA diving (Haddock & Heine, 2005) from a small vessel off the coast of Kailua-Kona (Hawai’i Big Island, 19°42’38.7" N 156°06’15.8" W), over 2000 m of offshore water. Some dives were diurnal, where we collected most of the specimens of *Iasis cylindrica*, *Cyclosalpa affinis*, *Cyclosalpa sewelli*, and *Brooksia rostrata.* We observed and collected most specimens of other species during night dives (blackwater diving). We recorded in situ underwater videos of salp colonies swimming using a variety of cameras including a stereo system (Sutherland et al. in review), a lightweight dual GoPro stereo system, a brightfield system (Colin et al. 2022), and a darkfield camera system.

### Measuring salp colony swimming speed

For most species, we collected and analyzed footage from multiple specimens (Table S1). We used a combination of spatially calibrated stereo video and 2D videos with a reference scale in the frame. From the stereo videos, we measured the XYZ positions of salp colonies in EventMeasure (SeaGIS). We implemented a cutoff in the RMS (root mean squared) point error estimate of < 2 mm. We complemented gaps in taxon sampling with archived 2D videos in the lab from previous expeditions to West Palm Beach (FL, USA) and the Pacific coast of Panama. For these 2D videos, we used the FFMPEG plugin in ImageJ to measure their XY positions in sequences where the colony was swimming horizontally within the focal plane. The colonies were assumed to be in the same plane as the scale bar so at same distance from the camera. However, in videos with a broad focal depth, this may not always had been the case, thus potentially introducing some measurement error. In addition, when loading the 2D videos in ImageJ, the virtual stack rendered a higher number of frames than those expected from the inherent frame rate. To address this, we calculated an operational frame rate for those videos dividing the number of ImageJ slices by the total duration.

We tracked the position of the first zooid’s viscera as well as the position of a reference particle in the water (methods described in Sutherland et al. in review) in 10-30 frames across 50-500 frame windows spanning 2-4s of swimming. The reference particle was a non-swimming organism (such as a foraminiferan or radiolarian) or a non-living particle. In addition, we recorded the pulsation rates of the specimens measured by counting the number of times the atrial siphon contracted in a known period. For each analyzed frame, we calculated the horizontal x, vertical y, and depth z (in the case of the stereo video measurement files) components of the relative positions of the frontal zooid as shown in Eq. 1.1-1.3.

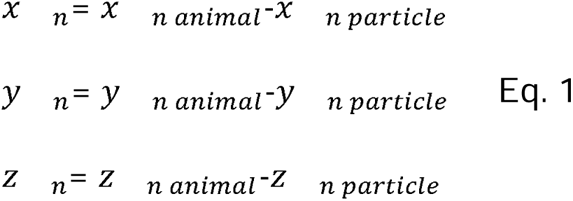

Then we calculated the instantaneous speeds using Eq. 2 (without the z component in the case of the 2D videos) given the known frame rate of each video.

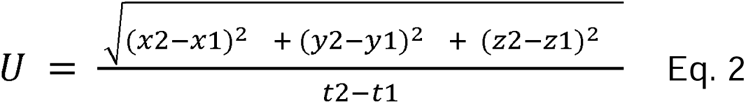

### Salp colonial architecture

To examine the relationships between locomotory efficiency and colonial architecture, we adopted the species-specific architecture characterizations and dorsoventral zooid rotation angle measurements for each species from Damian-Serrano et al. (2023). Using stills from the underwater videos, we measured zooid length, zooid width, and number of zooids in ImageJ. These measurements were repeated in at least three locations from each colony. When a distinct zooid size gradient was observed, we measured zooids in locations from the proximal, middle, and distal regions to capture the full range of variation in the specimen.

### Respiration measurements

We collected healthy, adult blastozooid (aggregate stage) colonies across 18 salp species (Table S2) during blue- and black-water SCUBA dives off the coast of Kona (Hawaii, USA) between September 2021 and May 2023. Specimens were sealed *in situ* with their surrounding water in plastic jars equipped with a Presens (Germany) oxygen sensor spot and a self-healing rubber port to allow for the injection of solutions without the introduction of air bubbles. We removed as many symbiotic animals from the salps as possible before closing the lid without damaging the colony. The same method was applied to one or more seawater controls to account for the oxygen demand of the local seawater’s microbiome. Several collection events occurred during each 20-60 min long SCUBA dive. Jars with larger animals were opened during the safety stop to allow them to re-oxygenate. Upon the divers’ return to the boat, we measured the initial oxygen concentration (mg/l) and temperature, and then repeated the measurements at intervals between 15min and 3h, for total periods ranging between 2h and 5h, depending on logistic constraints in the field and the rate of oxygen depletion. The exact interval time for each measurement was variable but recorded (Table S2).

To estimate the energetic expenditure of different salp species while actively swimming, we recorded the oxygen consumption of intact specimens while swimming inside the jar. To obtain a baseline of basal respiration rate (while not swimming), we anesthetized some specimens before the start of the first oxygen measurement time. A few specimens were used for paired experiments, where their swimming respiration was recorded for a few hours, then inoculated with the anesthetic, and recorded anesthetized for another set of hours. To anesthetize salps, we injected their jars with small volumes of concentrated (50 g/l) bicarbonate-buffered MS-222 through the rubber ports on the lids. We tailored the injection volume to the jar size aiming for a final concentration of 0.2g/l, following the methods in Trueman et al. (1984). We also injected some seawater control jars to evaluate the effect of MS-222 on oxygen concentration in seawater and found no effect.

When multiple seawater controls were collected using jars of different sizes, we paired each jar with the control that had the most similar volume. If among multiple controls only some were jars injected with anesthetic, we paired the anesthetized specimen jars with the injected controls and the intact specimen jars with the intact controls. In experiment 26 (see Table S2 for experiment numbers), the control jar was lost due to an encounter with an oceanic white tip shark, thus we paired those measurements with the nearest relative time points from the control jar in experiment 25, collected the same day hours earlier. At the end of each experiment, we identified the salp specimens used in the experiments to the species level, counted the number of zooids, and measured the zooid length (total length including projections) and the biovolume of the colony using a graduated cylinder. For those specimens where colony or zooid volume was not measured directly, we estimated the colony volume from their zooid length and the number of zooids using a Generalized Additive Model with the measured specimens.

We estimated the oxygen consumption rate for each specimen by fitting a linear regression of consumed oxygen mass (concentration by container volume) against the duration of the measurement series. We subtracted the slope calculated for the relevant control jar to the estimated slope of the animal jar. Since our seawater controls were not filtered, some experiments had abnormally high estimated background respiration rates, leading to negative values. We removed these data points before the analysis. To estimate biovolume-specific rates, we divided the rates by the colony volumes. We then compared the biovolume-specific respiration rates of active (swimming) and anesthetized specimens within each species, calculating the difference as a measure of biovolume-specific swimming cost respiration rate. We also calculated the relative investment in swimming as the proportion of biovolume-specific respiration rate comprised by the swimming-specific rate. To capture variability within species, we calculated the mean respiration rate of anesthetized specimens for each species and subtracted it from each specimen’s swimming-specific respiration rate within each species. We noticed that some species had higher average respiration rates among the anesthetized specimens than among the swimming specimens, leading to negative swimming-specific respiration estimates. We interpreted this anomaly as a systematic error due to the extremely low respiration rates of some species that fall within the effective detection limit of our experimental setup given the random variation range of respiration rates in seawater both in experimental jars and in control jars. Small absolute negative values get amplified into large relative values, especially in small animals with a minuscule biovolume denominator. Therefore, we removed the swimming specimens that had lower respiration rates than the mean anesthetized respiration rate for their species. We also removed two respirometry outliers of *Thalia* sp. which had extremely high swimming respiration rates (>7500 pgO2/ml/min, whereas all other measurements across species including other *Thalia* sp. were limited to 0-1700 pgO2/ml/min), which were likely due to amplification of experimental error (presence of organic matter or symbionts, underestimation of colony volume due to loss of tiny zooids in the sieves) with the small biovolume denominators in this species.

### Estimating costs of transport

We define the cost of transport (COT) as the amount of oxygen consumed per tissue volume per distance traveled by the colony. To estimate the COT, we divided the swimming-specific respiration rates by the mean swimming speed for each species measured from the stereo and 2D video data. Since the specimens used for speed measurements in the videos and those used in the respirometry experiments had different zooid sizes, we used the mean zooid-length per second speeds from the video measurements and then multiplied them by the actual zooid lengths of the respirometry specimens to estimate their absolute (mm/s) speeds. Pulsation rate estimates were taken from species averages from the video specimens. We also calculated the size-specific COT by transforming the swimming distances into zooid lengths measured from the respirometry specimens.

### Statistical Analyses

All data wrangling and statistics were carried out in R 3.6.3 (R Core Team 2021). To test for differences between architectures, we used two-sided t-tests. To test the relationships between pairs of continuous variables, we used linear models (as well as exponential models when comparing swimming speed to COT) and evaluated the significance of the slope parameter when compared against a flat slope. To evaluate the relative contribution of zooid size, pulsation rate, zooid number, and architecture type on swimming speed, we fitted a generalized linear model and evaluated the significance and proportion of variance explained by each factor using their partial R^2^.

## Results

Salp colony swimming speeds, pulsation rates, and respiration rates varied within and across species and colony architectures. Speeds measured with 2D methods were slightly slower than those measured with 3D methods within the species in which they overlapped. This is to be expected since 2D methods cannot account for the z (depth) component of the speed vector. When considering speed in terms of mm/s, we found no relationship between pulsation rate (effort) and absolute speed (p = 0.68, Fig. S1A), but a significant positive relationship with zooid-size corrected speed (p < 0.0001, Fig. S1B). Moreover, zooid length was positively correlated with speed, whether it is expressed as mm/s (p < 0.0001, Fig. S2A) or mm/pulse (p < 0.0001, Fig. S2B), in agreement with our initial hypotheses. Normalized swimming speeds (zooid lengths per pulse) allow for a more direct comparison of swimming speed across colonial architectures.

Salp species vary widely in their mean absolute colonial swimming speeds (Fig. 2A), with the slowest being under 6 mm/s in an oblique chain of *Thalia* sp., closely followed by the transversal chains of *Pegea confoederata* with 7.38 mm/s, and the fastest mean speed reaching 114 mm/s in the linear chains of *Salpa aspera*, closely followed by the linear chains of *Metcalfina hexagona* (107 mm/s) and *Soestia zonaria* (106 mm/s). The fastest individual specimen however, belonged to the linear chain species *Iasis cylindrica* with a speed of 176 mm/s. When correcting by zooid size and pulsation rate (Fig. 2B), *Soestia zonaria* was fastest with a mean velocity of 7.09 zooid lengths/pulse, followed by *Iasis cylindrica* (2.09 zooids/pulse) and *Salpa aspera* (2.03 zooids/pulse), whereas the oblique chains of *Thalia* sp. (0.37 zooids/pulse) were slowest, closely followed by the transversal chains of *Pegea* sp. with 0.43 zooids/pulse.

**Figure 2.**
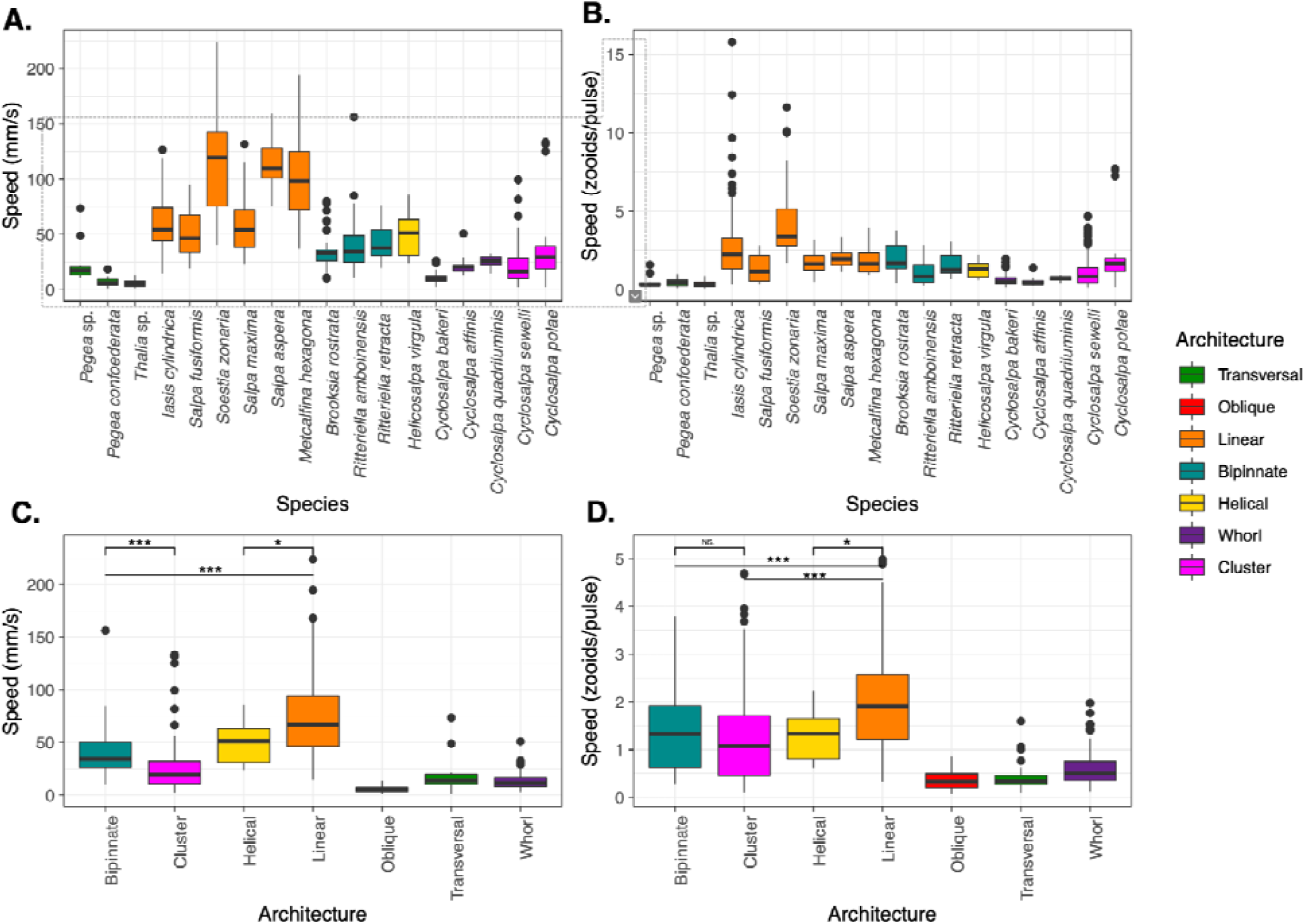
Boxplots showing the absolute (A) and corrected for body size and pulsation rate (B) swimming speeds recorded for each salp species and architecture (C, D) respectively. Colors correspond to colonial architecture types.

We find that swimming speed varies between colonial architecture types (Fig. 2C, D). Linear salp colonies are the faster than the rest (p < 0.001) with a mean speed of 73 mm/s (2.31 zooids/pulse), followed by helical chains (49.9 mm/s, 1.3 zooids/pulse), bipinnate colonies (39 mm/s, 1.41 zooids/pulse), and clusters (26.1 mm/s, 1.5 zooids/pulse). Among the slower architectures, we find whorls (13.4 mm/s, 0.64 zooids/pulse), transversal chains (16.7 mm/s, 0.44 zooids/pulse), and oblique chains (5.8 mm/s, 0.37 zooids/pulse). Our measurements of helical and oblique chains are limited to a single specimen, so this result should be interpreted with care.

Since linear architectures had faster mean swimming speeds (Fig. 2C, D), we investigated the relationship between swimming speeds with the dorsoventral zooid rotation angle, which represents the degree of linearity of the colony (Fig. 3). As hypothesized, species with more parallel (lower angles) dorsoventral zooid rotation present faster absolute speeds (-0.78, p < 0.0001) as well as somewhat faster size-and-effort corrected swimming speeds (-0.016 p < 0.0001). However, the latter relationship appears to be driven primarily by the distinctly fast relative speed of the perfectly linear (0° zooid rotation angle) *Soestia zonaria*.

**Figure 3.**
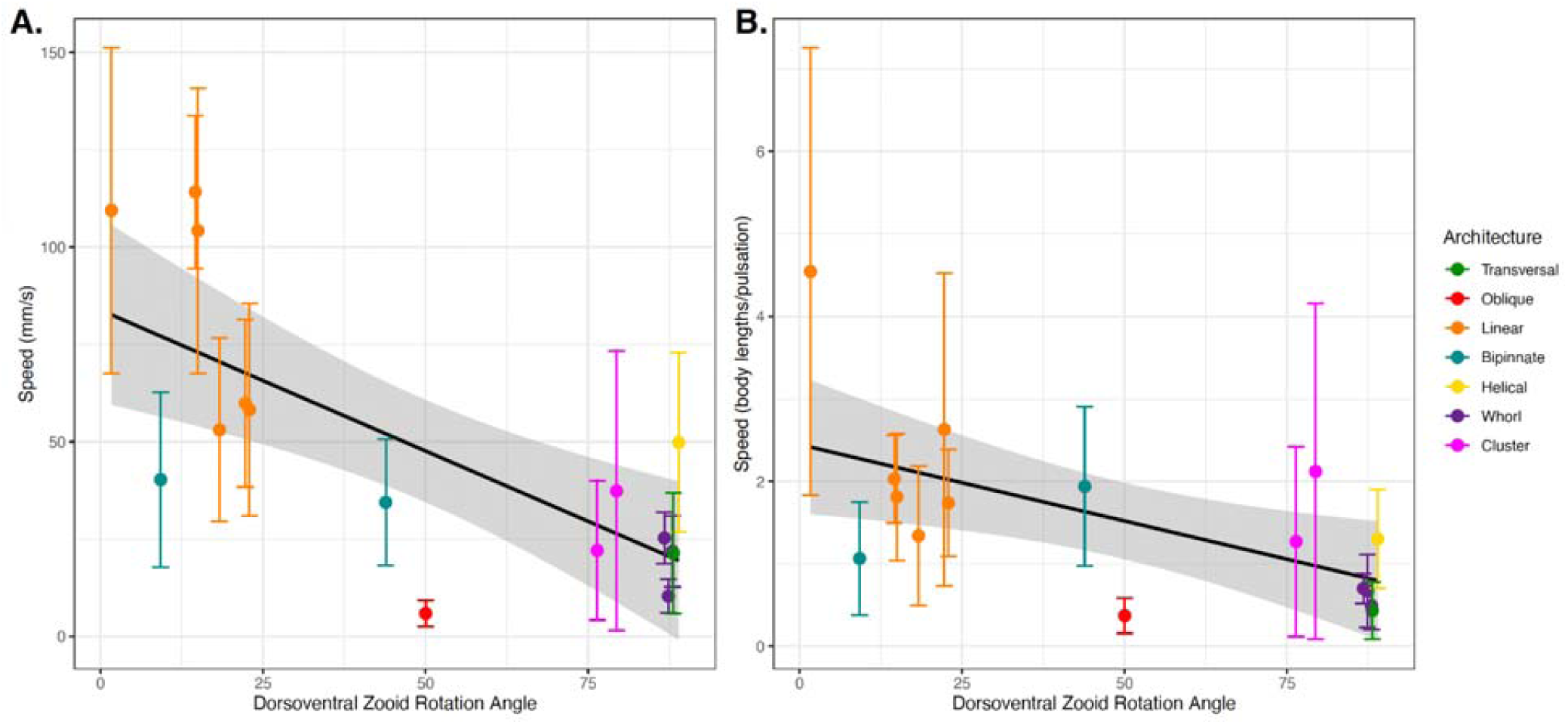
Absolute (A) and relative (B) colony swimming speed (specimen mean with standard errors) for each salp species across their degree of dorsoventral zooid rotation. Error bars indicate standard error. The color indicates colonial architecture. Gray areas indicate the 95% confidence interval of the linear regression (black line).

We compared how swimming speeds scale with the number of zooids in the colony (by species, see Fig. S3), and found differences between colonial architectures (Fig. S4). Swimming speed in whorls increased with number of zooids (0.08, p < 0.0001), which is incongruent with our frontal area hypothesis, but the data was limited to small numbers of zooids (4 to 13) and relatively slow speeds. As expected, linear chain architectures did increase in relative speed with the number of zooids (p < 0.001), as did bipinnate chains (p < 0.02), congruent with our frontal area hypothesis.

To further test our frontal area scaling hypothesis, we pooled the data from multiple architectures into scaling modes. We could then evaluate the overall relationship in colonies with a constant frontal area (linear, bipinnate, and helical species) and in colonies with scaling frontal area (transversal, whorl, cluster, and oblique species) with linear regression. This aggregation allowed the inclusion of data from architectures for which we only have one specimen (helical and oblique). When pooled by scaling mode (Fig. 4), the regression on colonies with a constant frontal area had a higher intercept on the swimming speed axis than in those with a scaling frontal area (1.54 and 1.09 zooids/pulse, respectively), reflecting the generally higher swimming speed of the former. Moreover, the regression on colonies with constant frontal area had a significant positive slope (0.02, p < 0.001), while the regression on those with scaling frontal area was not significant.

**Figure 4.**
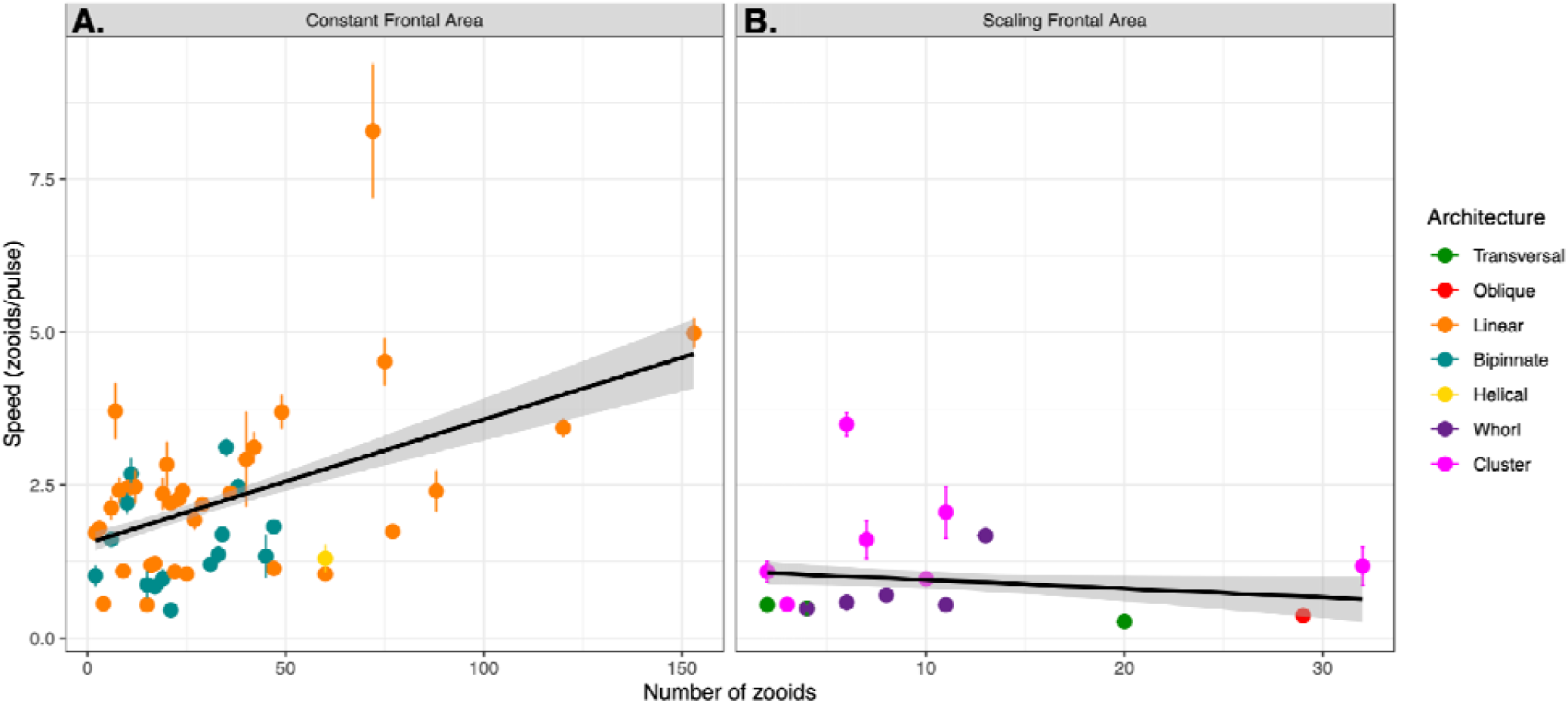
Linear relationships between relative swimming speed (zooid lengths per pulsation, specimen mean with standard errors) and number of zooids in the colony for constant (A) and scaling (B) frontal motion-orthogonal frontal area scaling modes. Gray areas represent the 95% confidence intervals of the regressions.

Putting together all the different organismal factors that we analyzed in this study, we calculated a generalized linear regression model to predict absolute salp swimming speed (U) from zooid length (L), pulsation rate (P), number of zooids (N), and colonial architecture represented as frontal area scaling mode (A) as expressed in Eq. 3. While our results suggest that the effect of N depends on A, we favored this simpler regression formula because it had a significantly lower (delta > 70) AIC score than those with interaction terms between N and A.

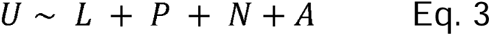

In this global model, we find significant effects (p < 0.001) from L, N, and A. We find that our global regression explains 36.76% of the variance in our swimming speed data: 5.78% is explained by zooid size, 3.52% by pulsation rate, 0.81% from zooid number, and 26.64% by the frontal scaling mode.

In addition to investigating the determinants of swimming speed in salp colonies, we also compared their respiratory physiology and the energetic efficiency of their swimming. The respiration rates of swimming and anesthetized salps in sealed jars at ambient temperature revealed broad differences between species (Fig. S5). Among swimming specimens (*Thalia* sp. high outliers removed), the highest mean swimming respiration rate was for *Ihlea punctata*, with an average of 1560.9 pgO2/min/ml. This species had also the largest difference between swimming and anesthetized respiration rates. The lowest mean swimming respiration rate (negative values removed) was for *Ritteriella amboinensis* with 33 pgO2/min/ml. The highest mean anesthetized respiration rate per ml of biovolume we recorded was for *Thalia* sp. with 715 pgO2/min/ml, while the lowest was for *Cyclosapa bakeri* (28 pgO2/min/ml).

We estimated the cost of transport (COT) as the biovolume-normalized metabolic cost of locomotion per unit of distance, as the mass of oxygen consumed both per mm traveled as well as normalized per body (zooid) length traveled. The species with the costliest locomotion was *Thalia* sp. (oblique architecture), followed by *Pegea* sp. (transversal architecture). We find that *S. zonaria* and *C. sewelli* have the lowest (most efficient) COT (Fig. 5A, B). Some of the differences between COT per mm and COT per zooid length are likely due to scaling with body size, as can be observed with the relative shift in the minuscule *Thalia* sp. (5.2 mm zooids) and the massive *Salpa maxima* (93.4 mm zooids). While linear architectures have the lowest mean COT values, these are not significantly lower than helical, bipinnate, whorls, or clusters (Fig. 5C, D). All these architectures have similar mean COT values that are much lower than those found in transversal and oblique architectures. These results do not support the hypothesis that more streamlined architectures have more energetically efficient locomotion.

**Figure 5.**
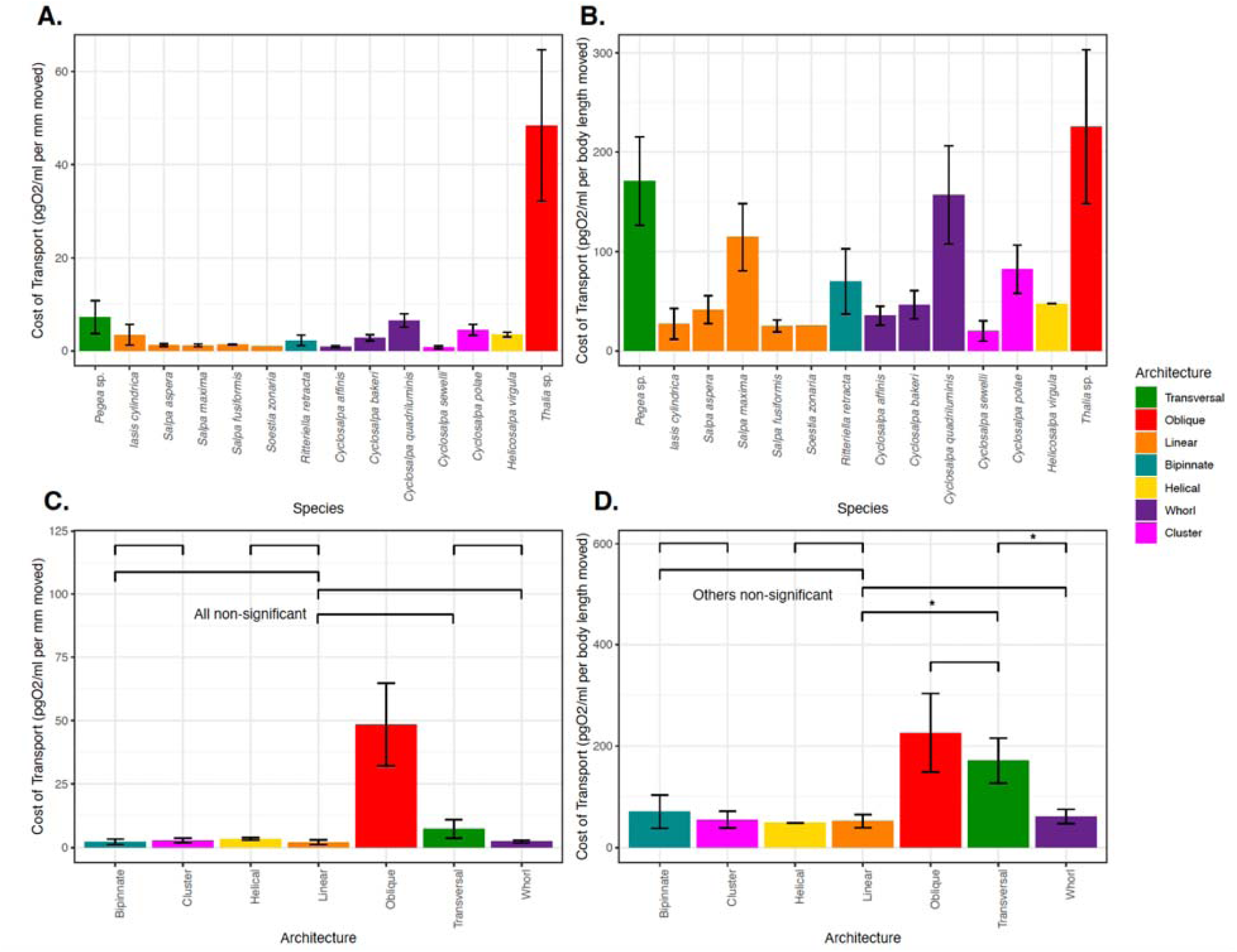
Mean cost-of-transport per mm (A) and per zooid length (B) moved for each salp species, and for each colonial architecture (C, D) with standard errors. Bar colors indicate colonial architecture.

When comparing the proportion of investment of metabolic costs into swimming (compared to the species mean baseline) across salp species (Fig. S6), eight species had locomotion budgets under 50%, and the other seven have budgets above 50%. The species with the highest relative investment in locomotion are *I. punctata* (94.5%) and *H*. *virgula* (83.1%), while the lowest investments were found in *C. affinis* (19.8%) and *Thalia* sp. (30.6%).

Upon noticing this variation, we examined whether observed effort (pulsation rate) scales with the measured proportion of energetic investment in swimming across species (Fig. S7) and found no significant relationship (p = 0.47) between them, and thus no support for the hypothesis that higher swimming effort incurs a higher metabolic effort.

We then compared the proportion of energetic investment in swimming to the COT values across species (Fig. S8) and found no relationship with absolute COT but found a positive relationship with zooid-length scaled COT (p < 0.001), indicating that species with more costly locomotion per zooid length invest a larger proportion of their energy budget in swimming. Finally, we compared the proportion of energetic investment in swimming with speed (Fig. S9). We found no relationship (neither in mm/s nor in zooids/s), indicating that faster swimmers do not invest more or less proportion of their energy budget into their locomotion efforts. We found that regardless of whether we consider transport in terms of absolute distances (Fig. 6A, linear regression p < 0.005, exponential regression p < 0.001) or relative to body lengths (Fig. 6B, linear regression p < 0.01, exponential regression p < 0.001), the COT decreases in species with faster swimming speeds.

**Figure 6.**
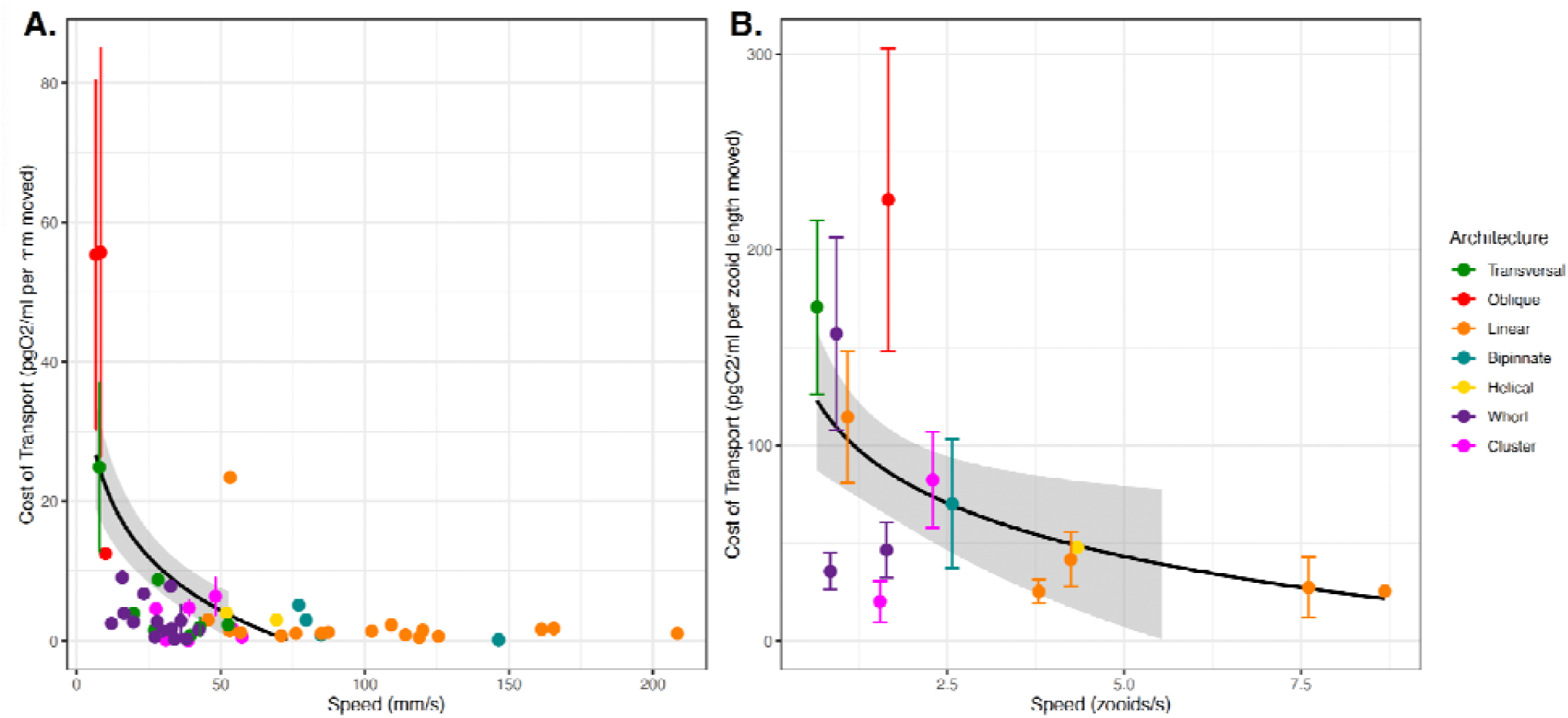
COT (specimen mean with standard error) per mm (A) and zooid length (B) moved across the specimen mean absolute (A) or relative (B) swimming speeds. The dot color indicates colonial architecture. Gray areas represent the 95% confidence intervals of the exponential regressions (black lines).

## Discussion

We compared the swimming speeds and costs of transport of salp colonies across the most comprehensive representation of salp species diversity. Our results show a wide range of colonial swimming speeds across salp species and architectures. Moreover, this study shows for the first time how salp colonial swimming speed scales with the number of zooids in the colony, suggesting that incremental propulsive power from additional zooids does not always produce higher swimming speeds.

### Architectural determinants of salp swimming speed

Colonial architecture was the strongest predictor of swimming speed, and though there is a large amount of unexplained variation which may relate to species-specific differences, behavioral, or environmental factors (see global GLM results). When ranking the different architectures by their swimming speed, our results are partially in agreement with our hypothesized ordination insofar as linear, bipinnate, and helical chains are the fastest, with transversal chains and whorls ranking lower. However, cluster architectures were faster than we anticipated, and oblique chains much slower than we expected.

We expected that swimming speed in colonial salps would be predicted by pulsation rate as a measure of swimming effort. Our results indicate that this relationship only exists when accounting for zooid size, suggesting an underlying relationship between pulsation rate and zooid size that may be masking its predictive power over absolute speeds. This is consistent with the distribution of our data and our observations in the field where larger salps pulsate at a slower rate than smaller ones. While Madin (1990) found no relationship between zooid size and speed in single zooids, we do find a significant increase in speed with larger zooid sizes, indicating that multi-jet propelled animals follow more similar scaling rules to vertebrate swimmers (Vogel 2008) than to single-jet propellers.

The relationship between the number of zooids and speed in linear chains is weaker than we would expect. This may be partly explained by the phenomenology behind more and less populous colonies. Salp colonies start their free-living phase when the developing buds detach from the solitary oozooid. This is when the colony is expected to have the maximum number of zooids since the zooid number only gets reduced as the colony splits or loses zooids to turbulence, disease, or predation. Therefore, colonies with higher numbers of zooids are typically composed of smaller, younger zooids. In linear architectures, these younger colonies could still be developing their dorsoventral rotation (Damian-Serrano & Sutherland 2023), thus effectively being more similar to oblique architecture. A less acute dorsoventral rotation angle would explain why these more numerous linear chains are not as fast as we would expect, given that our results support a significant relationship between this angle and swimming speed (Fig. 3).

Linear chains swam faster than all other architectures, including those that share a constant frontal area. One potential explanation for this difference could come from the relative thrust provided by the jets. Linear chains eject their jet plumes at very small angles (near parallel) to the axis of locomotion (Sutherland et al. in review). Bipinnate and helical chains (both with constant frontal area) have the atrial siphons (point of jet ejection) of their constituent blastozooids oriented at a wider angle (Madin 1990), which may lead to wider angles of their jets relative to the axis of locomotion. This in turn would result in a larger proportion of the force exerted by the jet to be applied as torque rather than thrust onto the colony. This hypothesis could be tested by measuring the 3D angles of the actual jets instead of the angles of the zooids since salps can use their atrial muscles and siphon morphology to direct the angle of their jets.

Finding that clusters can swim at speeds comparable to those of bipinnate and helical chains, even faster than whorls, defies our intuitive understanding of the mechanical properties of these colonies and thus warrants further investigation into how these species coordinate their jets to produce forward thrust. While oblique chains are architectural intermediates between transversal and linear chains, our results indicate that oblique chains are the slowest swimmers among salps. This incongruence may be explained by the fact that we only had speed data from one specimen (of *Thalia* sp.) with very small zooid sizes. Small salps might operate at notably lower Reynolds numbers than large ones, which may require a non-linear size correction for meaningful speed comparisons.

### Salp swimming speed and diel vertical migration

Salps are important players in the oceanic carbon cycle, grazing upon both phytoplanktonand bacteria (Henschke et al. 2016). Their carcasses and fecal pellets export large quantities of fixed carbon into the deep sea, accelerating carbon sequestration in the biological carbon pump (Wiebe et al. 1979, Décima et al. 2023). Part of this process is enhanced by the diel vertical migrations by some salp species though the distribution of this behavior across species diversity is poorly known. Off Bermuda, Madin et al. (1996) reported *Pegea* spp., *B. rostrata*, and *C. polae* as non-migratory, all of which we found to have slow swimming speeds. Other slow-swimmer species like *C. affinis* were found to only migrate a few meters through the diel cycle. The species *S. aspera, S. fusiformis*, *S. zonaria, I. punctata,* and *R. retracta* have been observed vertically migrating off Bermuda (Madin et al 1996, Stone & Steinberg 2014), which is congruent with our observations during fieldwork. These species all have constant frontal area and fast swimming speeds.

Vertical migrators need to be fast enough to follow the dark isolumes as they shift during dawn and dusk in time to maximize their exploitation of the food resources near the surface while avoiding exposure to daylight. Thus, absolute speed is important to the autoecology of these animals. Other *Salpa* species have also been reported as strong vertical migrators throughout the literature (Henschke et al. 2021, Madin et al. 2006, Pascual et al. 2017). A species that does not fit this pattern is *I. cylindrica*, a fast-swimming non-migratory species that spends night and day near the surface (Madin et al 1996; and pers. obs.). However, other studies do report moderate diel vertical migration for this species (Stone & Steinberg 2014), so it may be adapted for facultative vertical migration under specific oceanographic conditions. Some migratory species, such as *S. aspera*, are known to travel distances of over 800m at dawn and dusk, at rates predicted to require 5-10 m/min (83-166 mm/s) based on MOCNESS trawl intervals (Wiebe et al. 1979). These predictions are consistent with the speeds we recorded for this species (88-145 mm/s) and similar congenerics.

### Ecophysiological implications

While the importance of a few well-studied linear chain salp species in the biological carbon pump has been delineated, the question of whether this ecological role is generalizable to other salp species remains unanswered. In addition to vertical migration behavior, another likely important factor in their carbon flow is their respiration rate. The higher their respiration rate, the larger the proportion of assimilated carbon that will be released back into the water as dissolved carbon dioxide. This study provides the broadest taxonomic perspective on respiration rates (18 species, Fig. S5) and swimming cost of transport (14 species), finding 17-fold differences in their respiration rates and over 77-fold differences in their mean COT. Except for a few species with extremely high and low values, most respiration rates are centered between 0.2 and 1 µmol/g/hour, assuming a salp tissue density of 1.025 g/ml. In general, the respiration rates we estimated for salps are within the range of those reported in the literature (Trueblood 2019, Iguchi and Ikeda 2004). Compared to the metabolic rates estimated for the broader diversity of marine pelagic animals (Seibel & Drazen 2007), the rates that we measured for salps are in a similar range to those measured for *Salpa thompsoni* (Iguchi and Ikeda 2004). Our values are also similar to those measured by Seibel & Drazen (2007) in nemerteans, chaetognaths, and most fishes (0.1-1 µmolO2/g/h), which are generally higher than other gelatinous animals like ctenophores or scyphomedusae (0.01-0.1 µmolO2/g/h), but generally lower than those of cephalopods, crustaceans, or large fish (1-10 µmolO2/g/h). Salp species known to have strong vertical migration behaviors (*Salpa* spp., *S. zonaria, I. punctata,* and *R. retracta)* have low basal metabolic rates (Fig. S5) and low costs of transport. These results indicate that many non-migratory species, while likely still being important players in the biological carbon pump via their fecal pellet production, are releasing more of the consumed carbon as carbon dioxide near the surface than their more metabolically efficient relatives. The ultimate ecological outcome of each species needs to be assessed holistically, considering their microbial filtration and pellet deposition rate as well as their relative abundance in the water column.

Our metabolically calculated costs of transport range between 5-50 J/kg/m when converting the mg of oxygen to J via aerobic respiration free energy equations at 23°C. Our values are higher than the highly efficient 1-2 J/kg/m reported for salps in the literature (Bone & Trueman 1983, Gemmell et al. 2021), rather approaching the less-efficient values found in single jet-propelled invertebrates like scallops or squids. We suspect that COT calculated from mechanical parameters such as the displacement of water mass is not directly comparable to the COT calculated from respiration rates. Furthermore, we hypothesize that the standard aerobic respiration free-energy equation based on glucose is not an exact representation of the metabolic energy-conversion processes in salps, which may rely on a combination of sugars and fatty acids derived from their microscopic prey. Our results show that faster swimming species have lower COT (Fig. 6), which suggests that faster speeds and higher locomotory efficiency have a common cause, congruent with the hypothesis that both speed and efficiency depend on frontal drag forces. However, this hypothesis is not supported by the distribution of COT across architectures (Fig 5C, D), where except for oblique and transversal chains, all architectures present similarly efficient COT values. These results may be explained by the fact that swimming speed is an inversely proportional factor in the calculation of COT from respiration rates. Therefore, where we found surprisingly high and low speeds for clusters and oblique chains, we found surprisingly low and high COT values respectively. Perhaps there are other underlying explanatory factors linking swimming speed and swimming efficiency, such as muscle content, jet coordination, or jetting angles (thrust-to-torque ratios).

### Evolutionary implications

Across the evolutionary history of salps, linear chains have evolved multiple times independently from oblique ancestors (Damian-Serrano et al. 2023), suggesting the adaptive role of this architecture as a functional trait. Our results show that going from an oblique form to a linear one may confer significant advantages in locomotory speed and energetic efficiency. However, multiple colonial architectures, which we find to be slower swimmers (such as transversal chains, helical chains, whorls, and clusters) had also evolved from linear and oblique forms. This is incongruent with a scenario where natural selection strongly favors locomotion efficiency across all ecological niches of salps. Therefore, we hypothesize that in some lineages, the evolution of colonial architecture may be driven by ecological trade-offs with other non-locomotory functions. Alternatively, we hypothesize that, in some of these lineages, locomotion at the colonial stage may not be important enough for selection to maintain these highly hydrodynamic forms, allowing for neutral evolutionary processes to produce a diversity of non-adaptive forms.

#### Insights for bioinspired underwater vehicle design

Pulsatile jet propulsion is a promising avenue for bioinspired aquatic vehicles and robots (Mohensi 2006, Gohardini 2014, Yue et al. 2015). Multijet propulsion systems with multiple propellers akin to salp colonies are starting to be explored in an engineering context (Chao et al. 2017, Costello et al. 2015) with direct inspiration from gelatinous animals (Marut 2014, Krummel 2019, Bi et al 2022, Du Clos et al. 2022). Salp diversity provides a natural laboratory to explore the hydrodynamic implications of different multijet arrangement designs. Our findings underscore the importance of considering the scaling hydrodynamic properties of propeller arrangements to optimize speed and energy efficiency in bioinspired underwater vehicle design. While linear chain arrangements were the fastest and among the most energy efficient, robot (or vehicle) configurations such as a cluster form may confer unique object manipulation or maneuverability advantages. Our results show that these seemingly inefficient propeller configurations do not impose large disadvantages in terms of speed and fuel efficiency.

## Acknowledgments

We are grateful to the crew of Aquatic Life Divers, Kona Honu Divers for their assistance and support in hosting our offshore diving operations. We also wish to thank Marc Hughes, Jeff Milisen, Rebecca Gordon, Matt Connelly, Clint Collins, Paul Richardson, and Anne Thompson for their assistance during diving, collections, and filming operations in the field. Finally, we would like to thank Tiffany Bachtel for her valuable advice on the respirometry experiment design.

